# Lifelong Ulk1-mediated autophagy deficiency in muscle induces mitochondrial dysfunction and contractile weakness

**DOI:** 10.1101/2020.07.31.223412

**Authors:** Anna S. Nichenko, Jacob R. Sorensen, W. Michael Southern, Anita E. Qualls, Albino G. Schifino, Jennifer McFaline-Figueroa, Jamie E. Blum, Anna E. Thalacker-Mercer, Sarah M. Greising, Jarrod A. Call

## Abstract

The accumulation of damaged mitochondria due to insufficient autophagy has been implicated in the pathophysiology of sarcopenia resulting in reduced contractile and metabolic function. Ulk1 is an autophagy-related kinase that initiates autophagosome assembly and may also play a role in autophagosome degradation (i.e., autophagy flux), but the contribution of Ulk1 to healthy muscle aging is unclear. We found that Ulk1 phosphorylation declines in both human and mouse muscle tissue with age, therefore the purpose of this study was to investigate the role of Ulk1-mediated autophagy in skeletal muscle aging. At age 22 months (80% survival rate), muscle contractile and metabolic function were assessed using electrophysiology in muscle specific Ulk1 knockout mice (MKO) and their littermate controls (LM). Specific peak-isometric torque of the ankle dorsiflexors (normalized by tibialis anterior muscle cross-sectional area) and specific force of the fast-twitch extensor digitorum longus muscles were reduced in MKO mice compared to LM mice (p<0.03). Permeabilized muscle fibers from MKO mice had greater mitochondrial content, yet lower mitochondrial oxygen consumption and greater reactive oxygen species production compared to fibers from LM mice (p≤ 0.04). Altered neuromuscular junction innervation patterns and changes in autophagosome numbers and/or flux in muscles from MKO may have contributed to decrements in contractile and metabolic function. Results from this study support an important role of Ulk1-mediated autophagy in skeletal muscle with age, reflecting Ulk1’s dual role in maintaining mitochondrial integrity through autophagosome assembly and degradation. A lifetime of insufficient Ulk-1-mediated autophagy in skeletal muscle exacerbates age-related contractile and metabolic dysfunction.

## Introduction

Autophagy is an evolutionarily conserved cellular process for degrading damaged and dysfunctional proteins and organelles and has been strongly associated with the pathophysiology of sarcopenia (age-related reduction in skeletal muscle mass and function) ^1–3^. Studies investigating autophagy in older adults and rodents report less basal autophagy signaling as well as less mitochondrial-specific autophagy (i.e. mitophagy), signaling with age ^4–8^. Moreover, autophagy-related protein knockout models indicate an exacerbated aging phenotype supporting an important role for autophagy in healthy skeletal muscle aging ^7,9–12^. For example, Carnio et al. discovered geriatric mice with deficient ATG7, an autophagy-related protein important for autophagosome assembly, have weaker muscles that present with greater levels of muscle fiber atrophy and dysfunctional neuromuscular junctions (NMJs) compared to age-matched controls^7^. Autophagy is a dynamic process that can be altered not only by changing the number of autophagosomes assembled but also by changing the rate of autophagosome degradation (i.e., autophagy flux). In addition to changes in autophagosome assembly there is compelling research to suggest that autophagic flux is also decreased with age ^10–12^. Therefore, it appears that autophagy is inextricably linked with aging and reductions in both autophagosome number and flux may lead to an accumulation of damaged organelles and proteins which contribute to sarcopenia.

The accumulation of damaged and dysfunctional mitochondria with age is thought to contribute to sarcopenia due to metabolic insufficiency and a greater production of reactive oxygen species (ROS) ^13–16^. Damaged and ROS-producing mitochondria can be degraded through mitophagy, which is regulated in part through Unc-51 like autophagy activating kinase, Ulk1 ^17–19^. Ulk1 has long been thought of as an autophagy initiating protein that controls autophagosome assembly, but recent mechanistic investigations in yeast have suggested that Ulk1 may also be involved in signaling for autophagosome-lysosome fusion which leads to autophagosome degradation (autophagy flux) ^20^. We recently reported that muscle injury-associated degradation of dysfunctional mitochondria, through Ulk1-mediated autophagy, is critical for the timely recovery of muscle function ^21^. Thus, Ulk1 appears to play a mechanistic role in autophagosome assembly and flux, yet the extent to which Ulk1 influences sarcopenia is unclear.

The purpose of this study was to investigate the effects of lifelong insufficient Ulk1-mediated autophagy on aging skeletal muscle phenotypes in an effort to understand the contribution of reduced autophagy to sarcopenia. We hypothesized that lifelong Ulk1-defiency will exacerbate sarcopenia as indicated by a worsening of skeletal muscle contractile and metabolic function. Herein, we utilize highly sensitive electrophysiological techniques to assess muscle contractile and metabolic function in aged mice with genetic deletion of Ulk1 and their littermate controls.

## Methods

### Human Muscle Tissue Ethical Approval and Collection

All research involving human participants was approved by the Cornell University institutional review board and complies with the Helsinki declaration. Young (aged 20-40 y) and old (aged 60-80 y), male and female biopsy donors were recruited from Tompkins County, NY and the surrounding area. The following exclusion criteria were used: presence of a musculoskeletal disease, movement disorder, or other conditions known to influence skeletal muscle tissue (e.g., diabetes, cancer); high alcohol intake (>11 servings/wk for women, and >14 servings/wk for men); taking immunosuppressive or anti-coagulant medication; pregnant or breastfeeding; and recent weight changes or fluctuations. Biopsy tissue was collected following previously described methods ^22^, frozen in liquid nitrogen, and stored at −80°C until processing.

### Protein Isolation and Immunoblotting of Human Lysates

A 20-30 mg section of biopsy tissue was pulverized (Bessman pulverizer) and homogenized (VWR homogenizer) in lysis buffer (50 mM Tris, 150 mM NaCl, 1 mM EDTA, 1% NP-40, 0.5% Na-deoxycholate, 0.1% SDS, protease inhibitors [Roche], phosphatase inhibitors [Roche]). Lysates were incubated for 15 minutes on ice and centrifuged twice at 15,000 x g for 20 minutes at 4°C to remove insoluble material. Protein quantity was determined using a BCA assay (Thermo Scientific). Lysates were then diluted in laemmli loading buffer, heated to 95°C for 5 minutes, and separated on a 10% SDS gel (BioRad). Proteins were transferred from the gel onto a PVDF membrane (Millipore) at 30 V for 17 hours at 4°C.

### Ethical Approval and Animal Model

Animal protocols were approved by the University of Georgia Animal Care and Use Committee under the national guidelines set by the Association for Assessment and Accreditation of Laboratory Animal Care (A2017 08-004, UGA Animal Welfare Assurance #A3437-01). Muscle-specific Ulk1 knock-out mice (MKO) with *myogenin-Cre* and LoxP flanked *Ulk1* and their *myogenin-Cre* negative littermates (LM) were bred in-house and housed 5 per cage in a temperature-controlled facility with a 12:12 hour light:dark cycle and aged to either 16 months (middle-aged) or 22 months (old). 4 month old mice ^23^ were used for young adult controls in Ulk1 protein content analysis. All mice had *ab libitum* access to food and water throughout the duration of the experiment.

### Experimental Design

This study was originally designed to assess aging muscle health and function in a model of lifelong autophagy deficiency. The first cohort of mice consisted of Ulk1 MKO (n=10) and LM (n=10) mice that underwent longitudinal tracking of body mass and *in vivo* muscle strength every two months beginning at 12 months of age. Mice were sacrificed at 22 months of age based on a greater than 80% survivorship rate of our colony and published estimates ^24^. At sacrifice, muscles were harvested for *in vitro* muscle function analysis, mitochondrial function analysis, enzyme kinetics, immunohistochemistry, and immunoblots.

A second cohort of middle-aged mice (typically those aged 14-18 months) were used to assess autophagy related protein content and autophagy flux to determine how these might contribute to long-term physiological impairments. Mice were sacrificed at 16 months of age because the 16-month timepoint was the first longitudinal measurement where we observed an age-related decline in muscle function in the old mice.

A third cohort of young adult mice were sacrificed at 4 months old and the TA muscle was collected for immunoblots of Ulk1 and pUlk1(s555) protein content to compare to the middle-aged mice.

### In vivo assessment of muscle function

Peak isometric torque of the ankle dorsiflexors was assessed using a servomotor (Model 129 300C-LR; Aurora Scientific, Aurora, Ontario, Canada). Muscle torque (mN·m) was normalized to body mass (kg) to account for the large variability in body size. Detailed description of these methods is provided in the Supplementary Methods section.

### Glucose Tolerance Test

A glucose tolerance test was performed 4 days prior to sacrifice. Mice were fasted overnight (12 hours) before the test. On the morning of the GTT, a baseline measure of blood glucose was taken by tail snip bleed using a glucose monitor (EvenCare G3). Mice were then injected with sterilized D-glucose (200 mg/ml, i.p.) at 2 mg/g in normal saline and blood glucose was measured in 15 min increments for the next two hours.

### In vitro assessment of muscle function

Extensor digitorum longus (EDL) and soleus (SOL) muscles were excised and analyzed for force-generating capacities using a dual-mode muscle lever system (300B-LR; Aurora Scientific Inc., Aurora, ON, Canada). A detailed description of this method is provided in the Supplementary Methods section.

### Oxygen Consumption Rates

High resolution respirometry (Oroboros O2k) of permeabilized muscle fibers was used to assess mitochondrial oxygen consumption rates which were normalized to citrate synthase values to account for differences in mitochondrial content. A detailed description of these methods is provided in the Supplementary Methods section.

### ROS production

ROS production was assessed by quantifying resoflourin (red fluorescence) produced from the reaction of H_2_O_2_ and Amplex UltraRed (AmR, 10uM) catalyzed by horseradish peroxidase (HRP, 1U/ml) during the oxygen consumption measurements. ROS rates were normalized by oxygen consumption rates during leak respiration to account for differences in total oxygen flux through the system.

### Enzyme Assays

Citrate synthase enzyme activity was assessed to quantify mitochondrial content. The lateral portion of the gastrocnemius muscle remaining after fiber dissection was homogenized in 33mM phosphate buffer (pH 7.4) at a muscle to buffer ratio of 1:40 using a glass tissue grinder. Citrate Synthase activity was measured from the reduction of DTNB over time as previously described ^25^.

### Mitochondrial DNA analysis

Relative mtDNA copy number analysis was performed by the Oklahoma Nathan Shock Center on Aging by using the 2^−ΔΔct^ method as previously described ^26^.

### Chloroquine treatment (Autophagy flux measurement)

The second cohort of mice (middle-aged) were used to assess differences in basal autophagy flux between Ulk1 MKO and LM mice. Mice were i.p. injected with 100mg/kg of Chloroquine and sacrificed two hours after injection based on previously published guidelines for measuring autophagy flux ^27^. TA muscles were removed after sacrifice and processed for immunoblots of LC3II.

### Immunohistology

The TA muscle was used to assess fiber type differences and cross-sectional area (CSA) of fibers. The diaphragm muscle was used to assess neuromuscular junction integrity. A detailed account of the staining and imaging methods are provided in the Supplementary Methods section.

### Immunoblots

For autophagy-related protein content analysis, PVDF membranes were probed for the following proteins: Ulk1, pUlk1 (s555), DRP1, BNIP3, Pink1, Parkin, and LC3B. Primary antibody concentrations and a detailed description of the methods can be found in the Supplementary Methods section.

### Statistics

Changes in body mass, *in vivo* torque, EC50, and glucose tolerance were analyzed by two-way repeated measures analysis of variance (ANOVA) with main effects of genotype and time. Additionally, two-way ANOVA was used to analyze fiber type distribution and CSA by fiber type with main effect of genotype. Differences in *in vitro* force production, NMJ innervation, mitochondrial function, mitochondrial content, ROS production, mtDNA damage and autophagy protein expression between genotypes were analyzed by one-way ANOVAs with a main factor of genotype. Overall fiber CSA distribution was analyzed by Chi-squared. All data were required to pass normality (Shapiro-Wilk) and equal variance tests (Brown-Forsythe *F* test) before proceeding with the two-way repeated measures ANOVA. Significant interactions were tested with Tukey’s *post hoc* test using JMP statistical software (SAS, Cary, NC) to find differences between groups. Group main effects are reported where significant interactions were not observed. An α level of 0.05 was used for all analyses and all values are means ± SD.

## Results

Age-related declines in autophagy-related proteins LC3 (MAP1LC3A) and ATG7 have previously been reported, and these declines are thought to contribute to sarcopenia ^7^. To gain insight into changes in Ulk1 protein content with age, we analyzed total and activated Ulk1 (pSer555) protein content in old (60-80 y) and young (20-40 y) human muscle biopsy samples. There was significantly greater total Ulk1 protein content in old compared to young muscle (60%, p=0.0491), potentially reflecting the age-related loss of fast-twitch fibers and greater autophagy-related protein content in remaining slow-twitch fibers ^28^. Despite greater total Ulk1 protein, old muscles had 26% less activated Ulk1 normalized to total Ulk1 than young muscles, a marker of autophagy signaling (Fig 1A, p=0.0154). Similarly, middle-aged (16 months) mice contained 36% less activated Ulk1 normalized to total Ulk1 compared to young (4 month) mice (Fig 1B, p=0.0168).

**Figure 1:**
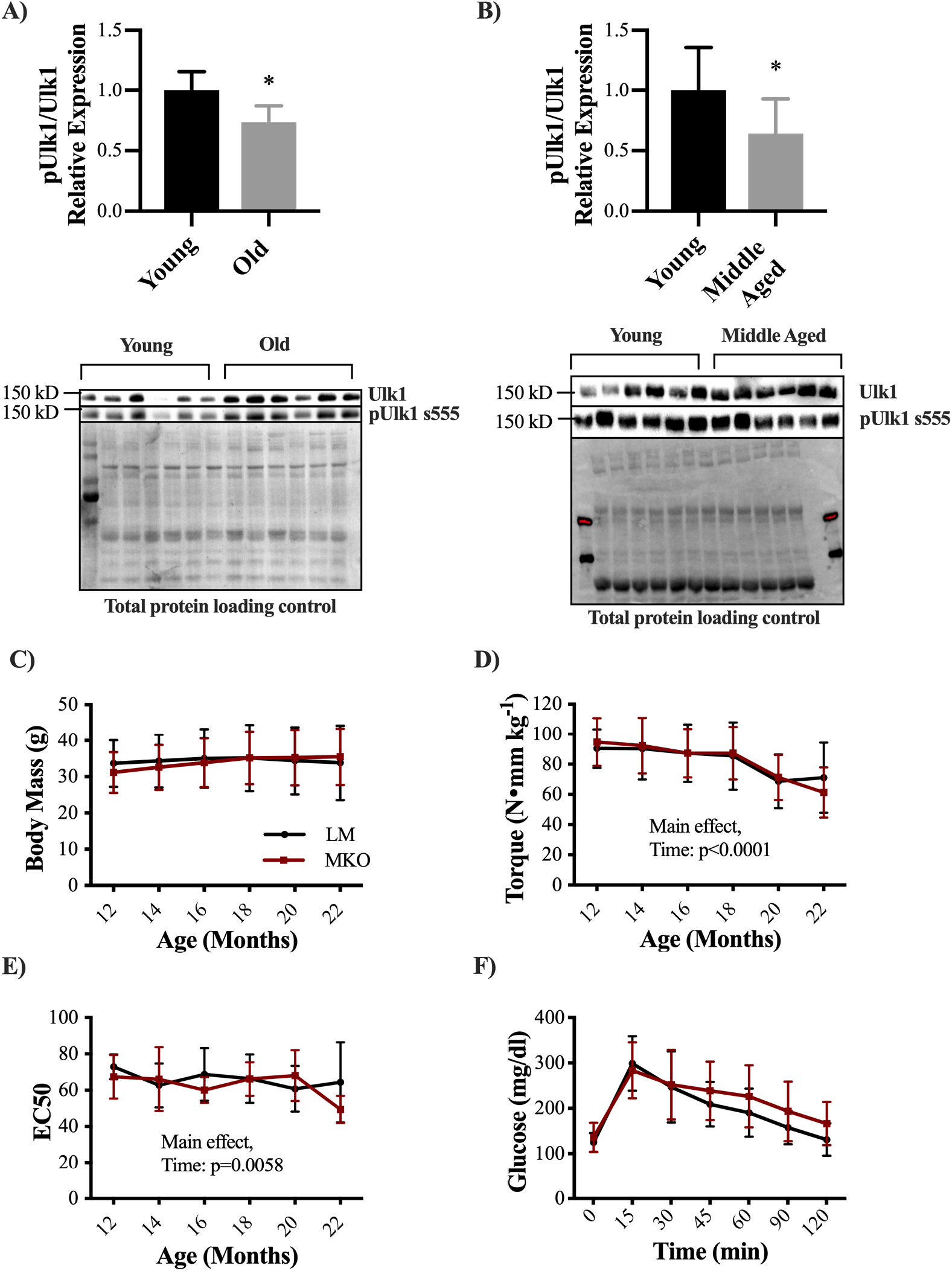
Decline in Ulk1 phosphorylation and skeletal muscle function with age. A) Relative expression of pUlk1 normalized to Ulk1 in young (20-40 y) and old (60-80 y) human muscle biopsy samples and representative immunoblot images. Mean ± SD n=6 samples. *= significantly different from young B) Relative expression of pUlk1 normalized to Ulk1 in young (4 months) and middle-aged (16 months) mice and representative immunoblot images. Mean ± SD n=7 mice. *= significantly different from young. C) Average longitudinal body mass measurements of Ulk1 MKO mice and LM controls starting at 12 months of age through 22 months. D) Average longitudinal *in vivo* muscle torque measurements from 12-22 months of age. E) EC50 calculated from longitudinal force frequency measurements. F) Average glucose levels recorded during a glucose tolerance test at 22 months of age in both Ulk1 MKO and LM control mice. All data are presented as mean ± SD n=10 mice. Main effects are listed where observed.

Ulk1 autophagy-deficient mice (MKO) at 22 months of age were used to interrogate how reduced Ulk1 protein signaling contributes to age-related skeletal muscle dysfunction. Body mass did not differ between LM age-matched controls and MKO mice or change significantly with time during the longitudinal portion of this study (12-22 months; Fig 1C, p=0.4181). We also investigated muscle masses after sacrifice and did not find any differences between genotypes in TA, EDL or soleus (SOL) muscle masses (Supp Fig. 1, p ≥0.07). However, gastrocnemius and heart muscle mass were significantly reduced in MKO mice when normalized to body mass (Supp Fig. 1, p≤0.04). Over the 12-month longitudinal period there was a 40% loss of peak isometric dorsiflexion torque independent of genotype (Fig 1D, p<0.0001). Sarcopenia is associated with a preferential loss of fast-twitch fibers, such that muscles take on a slow-twitch phenotype ^28^. We therefore analyzed the stimulation frequency at which dorsiflexor muscles reached half of their peak isometric torque in order to gain insight into potential fiber type shifts between LM and MKO mice (i.e., slow-twitch fiber summation is greater a lower stimulation frequencies). The frequency at which 50% of peak torque was reached (EC50) decreased by age, independent of genotype (70.1±9.9 vs. 52.4±20.3 Hz, 12- and 22-months, respectively, Fig 1E, Main effect: Age p=0.0058), indicating transition to a slow-twitch phenotype in the older mice. Prior to sacrifice, we used a glucose tolerance test to evaluate the impact of Ulk1 on insulin resistance in aged mice. MKO and LM mice showed similar glucose excursions and clearance after an injected glucose bolus (Fig 1F, p=0.8538).

To determine if muscle Ulk1 deficiency with age disproportionately affected fast-vs. slow-twitch muscles, *in vitro* contractility of the EDL and soleus muscles, respectively, was assessed in isolated muscles. We detected no difference in EDL maximal isometric force (Fig 2A, p=0.1750), however, when accounting for muscle CSA, EDL specific force production was 17% less in the MKO mice compared to the LM (Fig 2B, p=0.0277). No difference was detected between genotypes for either soleus maximal isometric force or specific force (Fig 2, p=0.2371), which suggests that fast-twitch muscles may be affected to a greater extent by Ulk1 deficiency throughout life.

**Figure 2:**
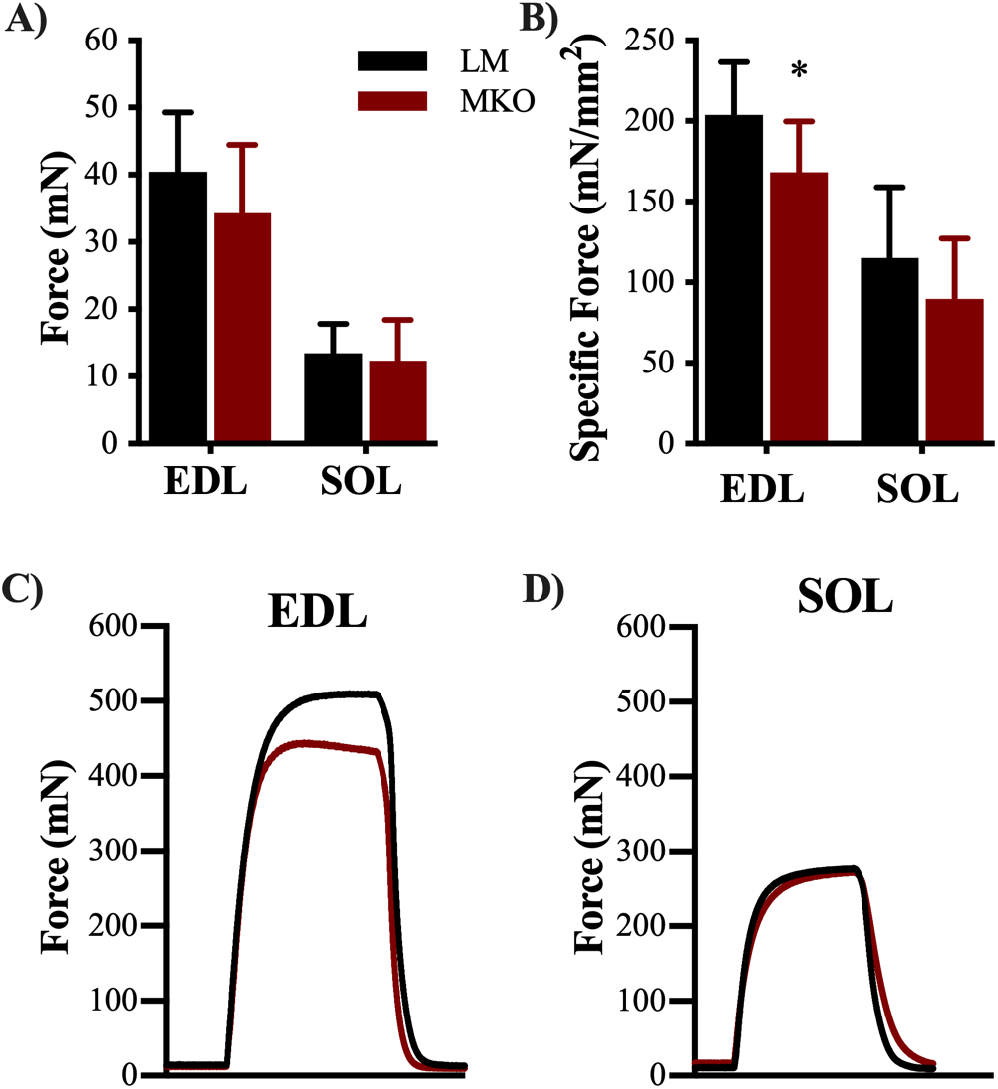
Ulk1 deficiency predominately affects type II muscle fibers strength. A) Average *in vitro* maximum contraction force from both EDL and SOL muscles (p>0.05) B) Average specific force from both EDL and SOL muscles (p=0.0277, and p=0.2371, respectfully). Representative torque tracing from C) EDL and D) SOL muscles. All data are presented as mean ± SD n=10 mice. *=Significantly different from LM

To determine the effects of Ulk1 on aging muscle anatomy, the primary ankle dorsiflexor muscle, the TA, was analyzed for differences in fiber number, fiber type specific distribution, and CSA. There was no difference in total fiber number between genotypes (2342±498 vs. 2036±678, LM and MKO, respectively, p=0.2640). There were expected differences in fiber type distribution of type IIa, IIx, and IIb fibers of the TA muscle (Fig. 3A, p=0.0210), however, there was no differences in the proportion of fiber types between genotypes. *In vivo* and *in vitro* contractile properties (e.g., twitch half-relaxation time) that can reflect fiber type shifts were also analyzed and there were no significant differences between genotypes (Supp table 1, p≥0.068). Expected differences in fiber type-specific CSA were observed, with IIa being the smallest and IIb the largest (Fig 3B, p<0.0001), yet no difference between genotypes was observed (p=0.9633). Intriguingly, the distribution of overall fiber CSAs independent of fiber type were shifted rightward so that the mean CSA of all fibers was 10% greater in the TA muscle of MKO compared to LM (1539±215 vs. 1692±248 μm^2^, LM and MKO, respectively) which is indicative of larger muscle fibers overall in MKO compared to LM (Fig. 3D, p=0.0035). The TA muscle contributes greater than 80% to peak-isometric torque of the ankle dorsiflexors, so we retroactively analyzed peak isometric torque normalized by the TA muscle CSA. Specific peak-isometric torque was 15% less in MKO mice compared to LM controls (Supp Fig 2, p=0.0078).

**Figure 3:**
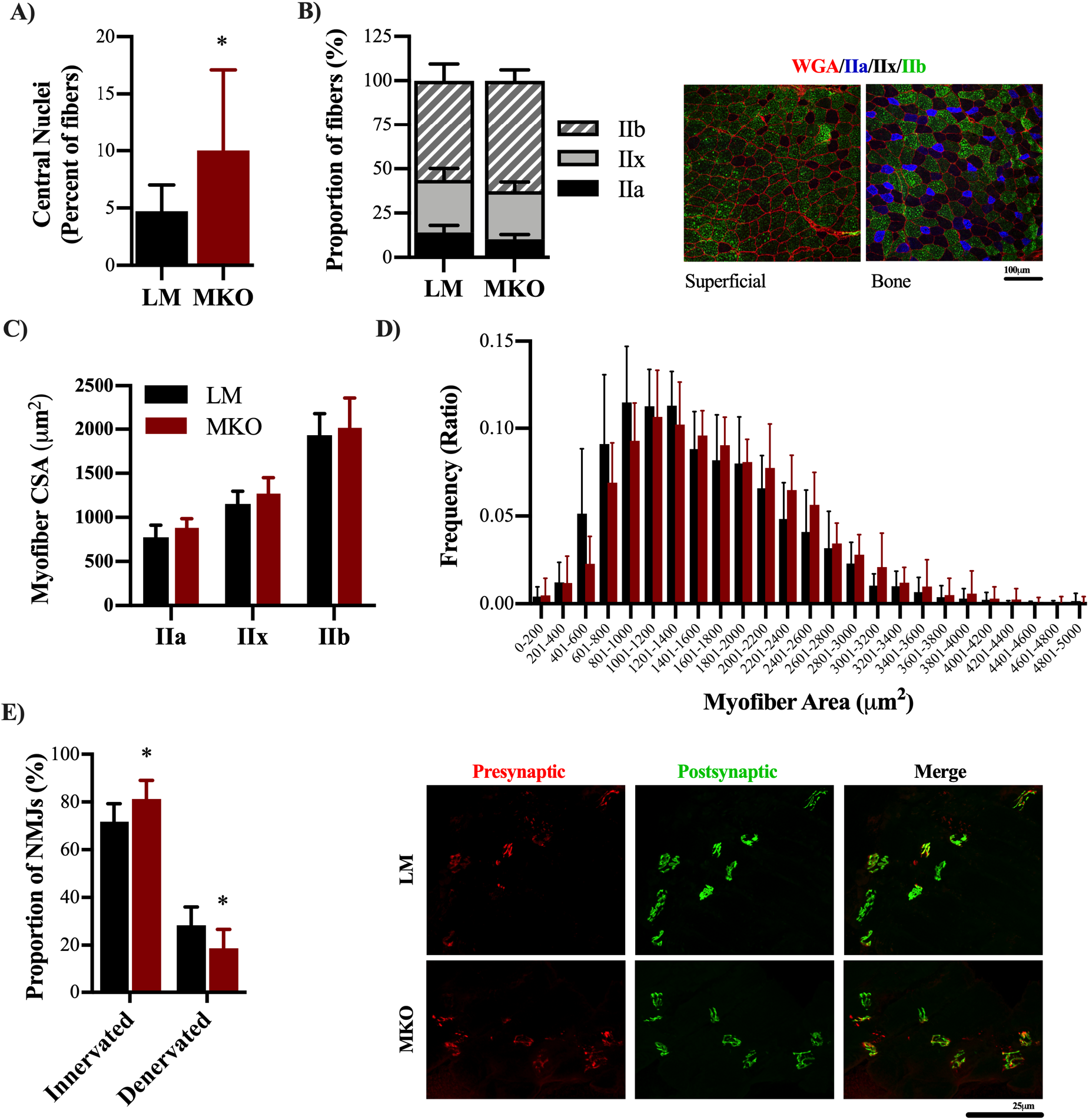
Ulk1 deficiency throughout life leads to enlarged but less functional muscle fibers and altered NMJs. A) Percent of centrally located nuclei (p=0.0487) B) Distribution of fiber types within TA muscle sections analyzed and representative images of fiber type staining. B) CSA of different fiber types D) Frequency distributions of fiber CSAs from TA muscles. (p=0.0035) E) Average number of innervated and denervated NMJs in the diaphragm of MKO and LM mice with representative image of neuromuscular junction staining (p=0.0227). All data are presented as mean ± SD. *= significantly different from LM

Centrally-located nuclei, an indication of muscle fiber death in the absence of injury, were assessed in TA muscles from LM and MKO mice. In agreement with a previous report involving muscle-specific Atg7 knockouts^7^, there was a greater percentage of centrally-located nuclei in autophagy deficient MKO muscles compared to LM (Fig. 3B, p=0.0487). Previous aging research and autophagy-deficient models have also revealed changes in NMJ integrity. ^7,29^ Therefore, to determine if Ulk1 influenced NMJs we assessed the innervation of diaphragm muscle NMJs between MKO and LM mice (Fig 3G). The diaphragm muscle displays all fiber types and is necessary for life, providing evaluation of a muscle in-between the predominantly slow- and fast-twitch limb muscles. Interestingly, the old MKO mice had a greater proportion of innervated fibers and less denervated fibers compared to age-matched LMs (Fig 3E, p=0.0227). The frequency of denervation in the age-matched LM is comparable to that previously reported ^29^, and follows a period of rapid denervation between middle and old age. We suspect our evaluation comes at a time in which many fibers have been lost to lack of innervation (i.e., there was survivor bias). Furthermore, we interpret this to indicate Ulk1 deficiency across the lifespan accelerated sarcopenia-related fiber death (i.e., centrally-located nuclei) and supports evaluation of skeletal muscle at an earlier age.

Ulk1-mediated autophagy is one of the main degradative pathways for skeletal muscle mitochondria, therefore we decided to investigate mitochondrial content, function, and ROS production after lifelong Ulk1 deficiency. Mitochondrial content of the gastrocnemius muscle (assessed through citrate synthase activity) was 37% greater in the MKO muscle fibers compared to LM (Fig. 4A, p=0.0443). Despite this, permeabilized gastrocnemius muscle fibers from MKO had 24% less State III (max physiological) and 25% less uncoupled (max) respiration compared to LM controls (Fig. 4B, p<0.02), which suggest impaired mitochondrial function. In agreement, MKO also produced more ROS per unit of oxygen flux than the LM permeabilized muscle fibers (Fig. 4C, p=0.0428). To determine the extent to which Ulk1 deficiency and greater ROS production damaged mitochondria, we assessed mitochondrial DNA content in the gastrocnemius muscle and found that it was decreased in MKO muscles compared to LM control muscles (Fig 4D, p=0.0309).

**Figure 4:**
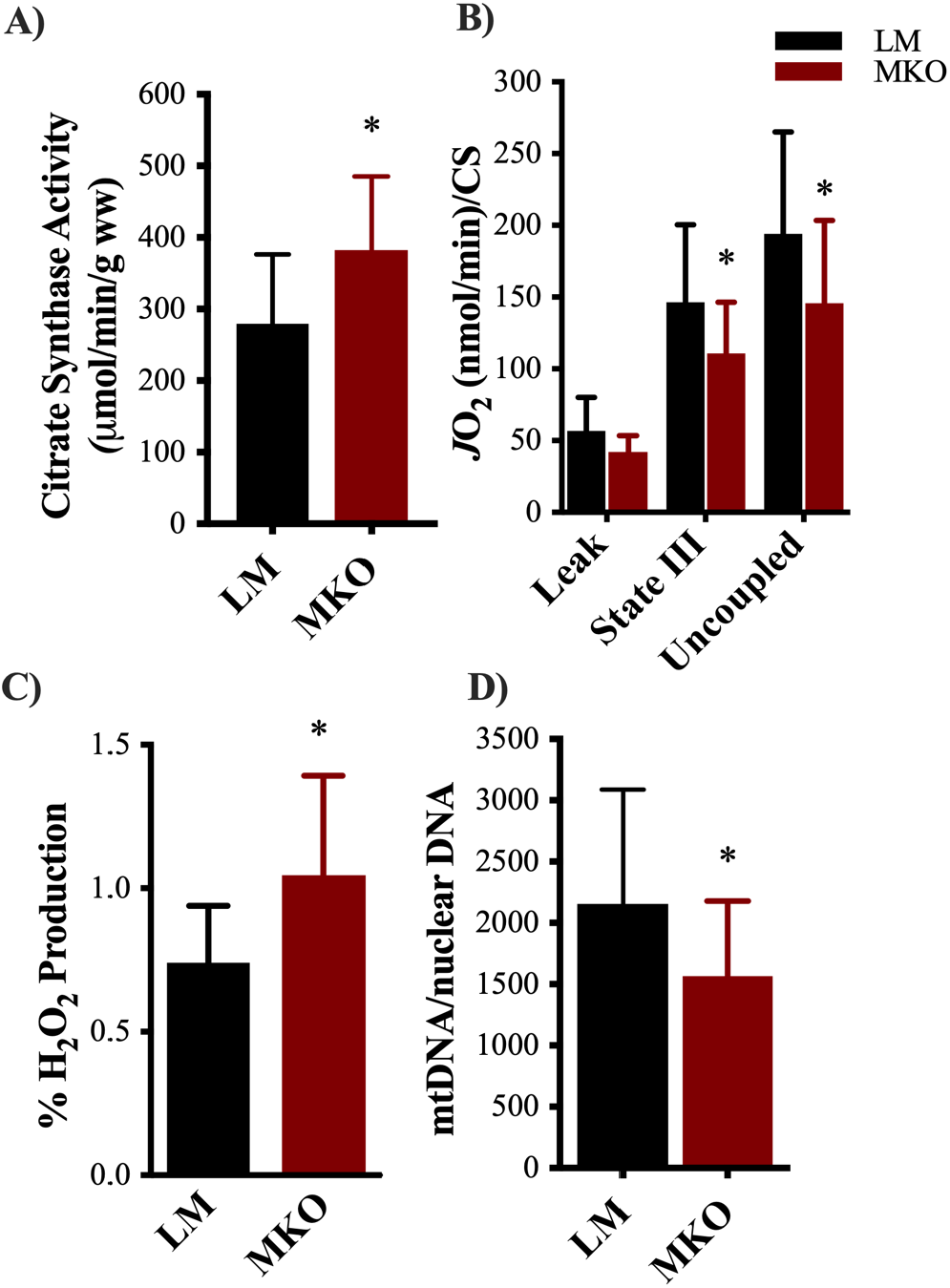
Lifelong Ulk1 deficiency results in an accumulation of dysfunctional mitochondria. A) Mitochondrial content assessed through citrate synthase enzyme kinetic rates (p=0.0443). B) Mitochondrial oxygen consumption during Leak, State III, and uncoupled respiration normalized to citrate synthase rate (p=0.0035 and p=0.0118, respectfully). C) Percent ROS production normalized to oxygen consumption during leak respiration to account for differences in oxygen flux (p=0.0428). D) Mitochondrial DNA content normalized to nuclear DNA content (p=0.0309).

We utilized a second cohort of middle-aged MKO and LM mice to better understand disruptions in autophagy signaling that may elicit the observed aging phenotypes. Middle-aged mice were selected based on the timepoint at which peak isometric torque significantly decreased in our old (22-month) cohort of mice. In a panel of proteins typically indicative of mitophagy signaling (DRP1, BNIP3, Pink1 and Parkin), there were no differences between genotypes in middle-aged mice (Fig. 5A, p>0.05). Basal autophagy flux was assessed by an acute treatment with the lysosomal inhibitor Chloroquine (CQ). CQ treatment resulted in greater accumulation of LC3II protein content independent of genotype (Fig. 5C, p=0.0007), and MKO mice had less LC3II protein content independent of treatment (Fig. 5C, p=0.0341). These data suggest that autophagosome number and/or degradation is impaired in MKO mice but is inconclusive on the extent to which autophagy flux is altered specifically.

**Figure 5:**
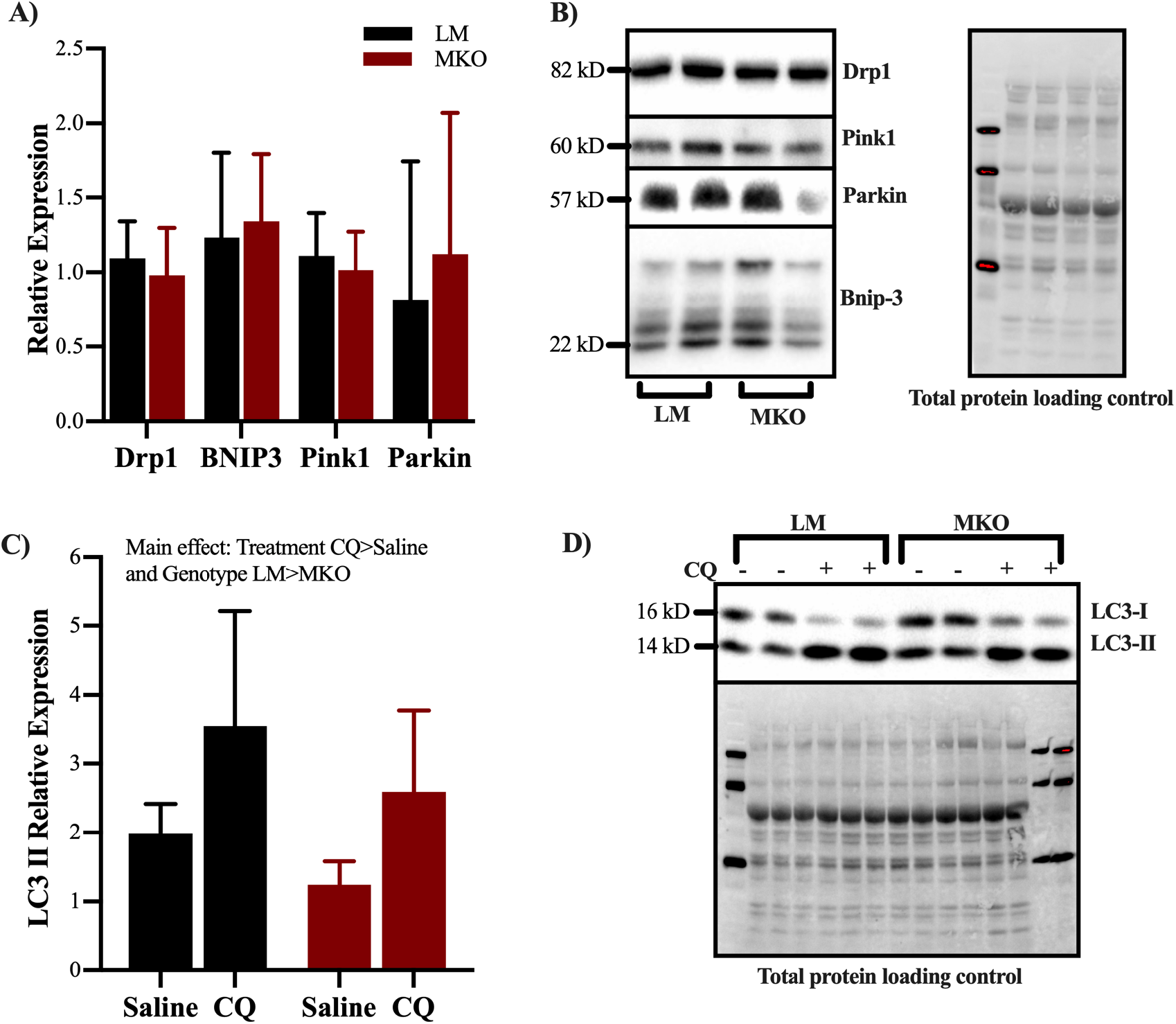
Mitophagy and autophagy flux in middle aged Ulk1 MKO mice. A) Relative expression of mitophagy related proteins DRP1, BNIP3, Pink1, and Parkin (p>0.05), B) Representative immunoblot panel C) Relative expression of LC3II in middle aged mice treated with Choloroquine or saline as a control (Main effect of Treatment and Genotype p=0.0007 and p=0.0341, respectfully). D) Representative immunoblots of LC3.

## Discussion

Herein we provide strong evidence in support of our hypothesis that lifelong Ulk1-defiency exacerbates sarcopenia, as indicated by a worsening of skeletal muscle contractile and metabolic function. Both LM and MKO mice exhibited expected age-related declines in muscle strength, yet muscle weakness was markedly greater in MKO mice when muscle CSA was accounted for in the EDL and TA muscles (Fig 2 and Supp Fig 2). Muscle fibers from MKO mice had poor mitochondrial respiration when normalized by mitochondrial content and produced more mitochondrial-derived ROS compared to muscle from LM mice, which supports the notion of greater mitochondrial dysfunction in MKO mice (Fig. 4). An accumulation of dysfunctional mitochondria could, in part, explain the advanced sarcopenia phenotype in MKO mice. Interestingly, we detected no difference in protein content related to mitochondrial removal (DRP1) or mitochondrial targeting for autophagosome encapsulation (BNIP3, Pink1, Parkin) between LM and MKO mice (Fig. 5). However, Ulk1 deficiency might have influenced autophagosome assembly and/or degradation as our chloroquine experiment showed less LC3II protein content in MKO muscle compared to LM (Fig. 5).

The decline of mitochondrial function throughout life is thought to be a major contributor to many aging phenotypes ^2,3,10,12^. We observed herein that with chronically deficient Ulk1, there is an exacerbated accumulation of damaged mitochondria by 22 months of age, which produce more ROS and have less mtDNA (a sign of damage) (Fig. 4). Similar findings of accumulated damaged mitochondria have been found in other aging autophagy knockout models ^7,9^. Interestingly, our data revealed through weaker EDL muscles that type II fibers were preferentially affected by deficient Ulk1 potentially through this accumulation of damaged mitochondria (Fig. 2). In support of this claim, Type II fibers have less endogenous antioxidants and less overall mitochondrial maintenance signaling ^28,30^. In other words, type II fibers may accumulate damaged mitochondria more quickly, produce more ROS as a consequence, and have less endogenous protection from ROS damage ^28,30^. Finding ways, such as exercise, to stimulate more mitochondrial maintenance signaling, particularly mitophagy, and endogenous antioxidant signaling with age may be a valuable therapeutic to preserve type II fiber function throughout life.

Ulk1 is one of the few autophagy-related kinases and its various roles in autophagy signaling are still being elucidated. Ulk1 appears to play a unique role in mitophagy signaling as it is sensitive to shifts in energy states and is post-translationally modified by energy sensing proteins like AMPK ^17,31^. AMPK phosphorylates Ulk1 at s555 prompting Ulk1 to co-localize to the mitochondria which is responsible for inducing mitophagy, as is the case in response to exercise ^19^. Specifically, AMPK-induced Ulk1 co-localization recruits’ autophagy-related machinery and lysosomes to the mitochondria to complete autophagic degradation ^19,20^. Not surprisingly, aged AMPK-deficient mice have weaker muscles with enlarged and damaged mitochondria (reduced mtDNA and increased ROS) similar to the data presented herein with Ulk1 MKO ^9^. Overall, this strongly supports the AMPK-Ulk1 signaling cascade for the degradation of damaged mitochondria though mitophagy is important with age. Because autophagy is a dynamic process, degradation and flux through the process must also be considered when assessing overall autophagic function. We have previously shown that although autophagy signaling is greatly upregulated in response to muscle injury, autophagic flux does not increase to the same extent which results in an autophagosome clearance bottleneck ^32^. It is unclear whether other scenarios result in a bottleneck, however, recent research has expanded the role of Ulk1 beyond autophagosome regulation and associated it with regulating autophagy flux ^20^. Wang et al. reported that in yeast, Atg1 (a Ulk1 homologue) kinase activity regulates the tethering of autophagosomal and lysosomal SNARE proteins which leads to autophagolysosome fusion and subsequent degradation ^20^. Post-translational modifications and kinase activities of Ulk1 were not within the scope of this paper, however, the absence of Ulk1 did not alter the expression of mitophagy proteins. This does not necessarily mean that Ulk1 is not critical for mitophagy proteins, it may simply reflect that the alternative autophagy pathways compensate ^19,33^.

Sarcopenia is associated with a reduction in motor units and some of the denervated muscle fibers can be inappropriately reinnervated such that a slow-twitch fiber is innervated by a fast-fatigable motor unit leading to altered recruitment patterns and changes in NMJ structure. Carnio et al. investigated the relationship between autophagy and NMJs specifically using ATG7 knockout mice and found that the muscle fibers from KO mice were expressing more NCAM, an attractant for alpha motoneurons, as a way to recruit new terminal axons to NMJs ^7^. It is unclear the extent to which our results support the work of Carnio et al., as our aged autophagy deficient mice actually appeared to have greater innervation (Fig. 3) ^7^. NMJ structure and innervation are a dynamic, ongoing processes and there is potential that we captured ongoing reinnervation in the MKO mice subsequent to loss of innervation. Alternatively, the absence of Ulk1 was associated with an undefined compensatory mechanism to increase alpha motoneuron innervation in order to try and improve muscle contractile function. Nonetheless, data presented here and reported by Carnio et al. are in agreement that deficient autophagy results in altered NMJ structure and innervation ratios ^7^, and autophagy may be a therapeutic target to mitigate NMJ changes with age.

Throughout life there is ongoing myonuclei turnover in uninjured muscle fibers in order to maintain muscle homeostasis. Specifically, satellite cells are reported to be responsible for this myonuclei turnover and this process results in increased centrally-located nuclei with age, a marker typically reflecting muscle injury and ongoing repair ^34–36^. It is unclear why centrally-located nuclei increase with age, but it appears to coincide with age-related changes in NMJ integrity ^35,37^. Autophagy deficiency may pre-dispose fibers to NMJ remodeling with age, and this could influence satellite cell dynamics and/or a muscle fibers susceptibility to contraction-induced injury based on changing motor unit recruitment patterns. The inverse is also possible, as we reported a protracted recovery process after injury in autophagy deficiency muscle and this could influence satellite cell behavior and NMJ integrity. Toward therapeutics, caloric restriction (a potent autophagy stimulus) decreases NMJ denervation and centrally located nuclei in aged mice ^37^. Collectively, there is support for further investigating the role of autophagy in mediating the relationship between NMJ and myonuclei maintenance in aged muscle.

The influence of sex and age on autophagy is an interesting conundrum that requires further investigation. A limitation herein is that we included both male and female mice in this study, but we were statistically underpowered to detect specific sex differences. There were three data outcomes that showed significant trends in sex differences, centrally-located nuclei were greater in MKO males, and in our mitophagy panel we detected that males had greater BNIP3 and decreased DRP1 protein expression independent of genotype. To our knowledge, there has been limited research on sex specific differences in autophagy. While one study linked estrogen receptor signaling to autophagy and mitochondrial function ^38^, whether testosterone signaling is involved in autophagy is unknown. Deciphering the sex specific differences in autophagy regulation with age may be an important avenue for further research.

In summary, lifelong Ulk1 deficiency results in an accumulation of dysfunctional, ROS producing mitochondria that coincides with reduced muscle force production, altered NMJs, and increased centrally-located nuclei. This aging phenotype may be due to a reduction in autophagosome formation and degradation as a consequence of absent Ulk1 signaling. Considering that Ulk1 activation naturally declines with age and its dual role in autophagosome formation and degradation, Ulk1 provides a strong potential therapeutic target to maintain muscle quality throughout life.

## Acknowledgements

We acknowledge Georgia Partners in Medicine REM seed grant (to JAC), and the Assistant Secretary of Defense for Health Affairs endorsed by the Department of Defense, through the Clinical & Rehabilitative Medicine Research Program, FY17 Neuromusculoskeletal Injuries Rehabilitation Research Award (W81XWH-18-1-0710 to JAC and SMG). Opinions, interpretations, conclusions and recommendations are those of the authors and are not necessarily endorsed by the Department of Defense. This material is based upon work supported by the National Science Foundation Graduate Research Fellowship Program under Grant No. DGE-1650441 (to JEB). Any opinions, findings, and conclusions or recommendations expressed in this material are those of the authors and do not necessarily reflect the views of the National Science Foundation.

---

**Supplemental Table 1:**
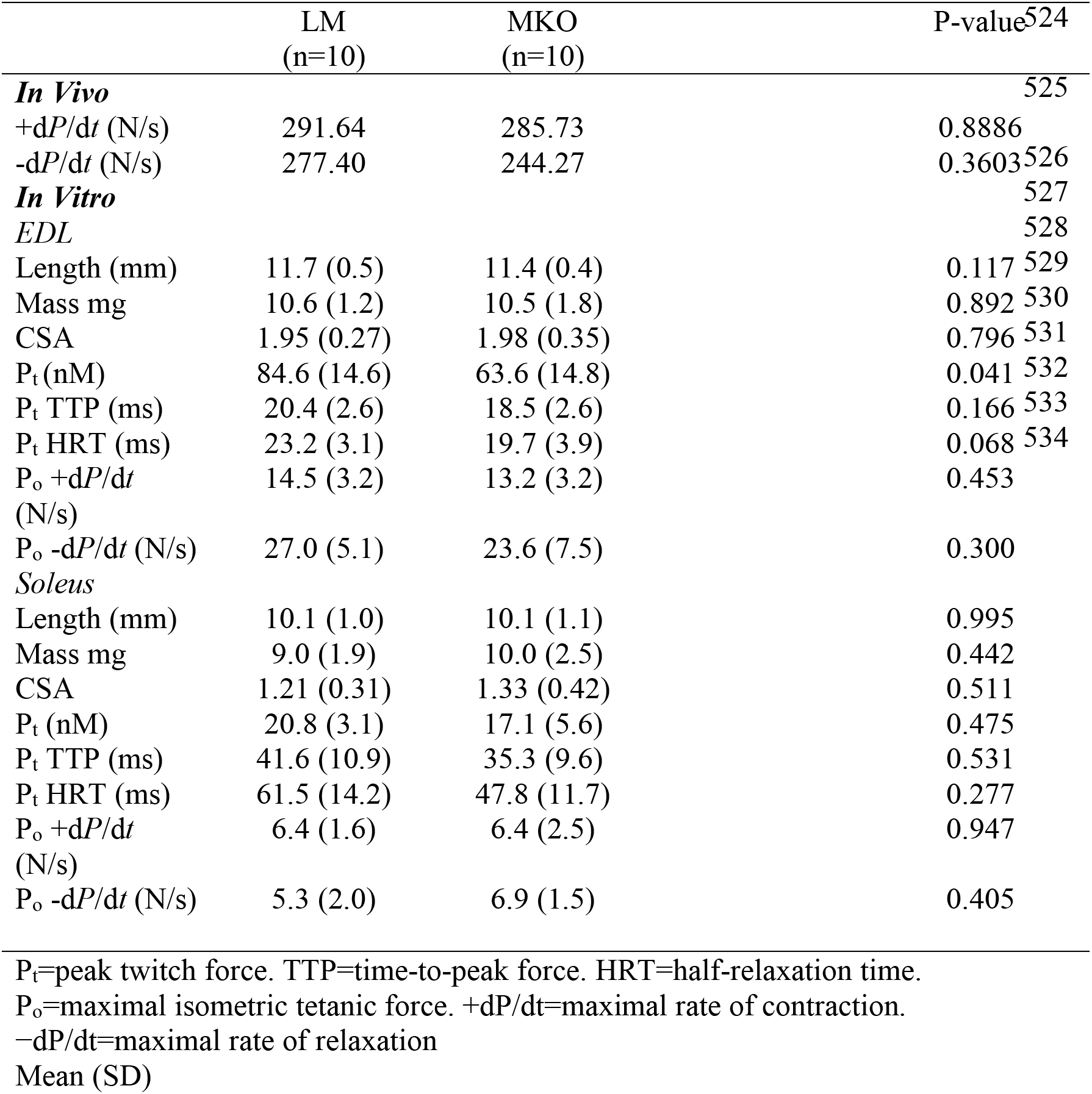
*in vivo* and *in vitro* contractile properties

## Supplemental Figures

**Supplemental Figure 1:**
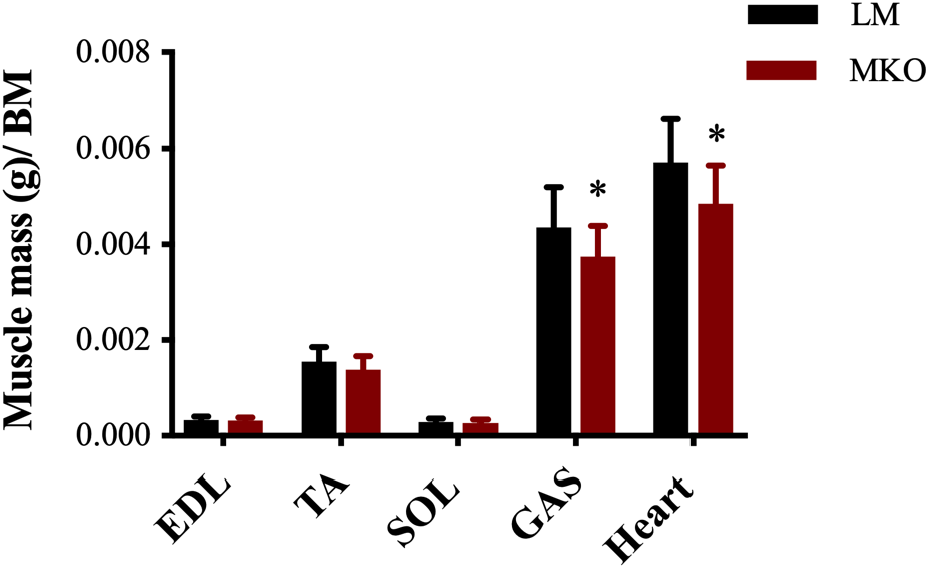
Muscle masses in old MKO and LM mice. Muscle masses normalized to body mass. All data are presented as mean ± SD n=10 mice. *= significantly different from LM

**Supplemental Figure 2:**
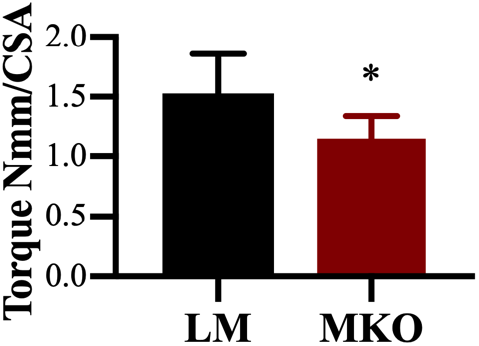
*in vivo* torque normalized by CSA in vivo torque normalized to CSA to account for fiber size differences between genotypes. All data are presented as mean ± SD n=10 mice. *= significantly different from LM.

## References

1. Brunk UT, Terman A. The mitochondrial-lysosomal axis theory of aging: accumulation of damaged mitochondria as a result of imperfect autophagocytosis. European journal of biochemistry. 2002;269(8):1996–2002.

2. Marzetti E, Calvani R, Cesari M, et al. Mitochondrial dysfunction and sarcopenia of aging: from signaling pathways to clinical trials. The international journal of biochemistry & cell biology. 2013;45(10):2288–2301.

3. Rajawat YS, Hilioti Z, Bossis I. Aging: central role for autophagy and the lysosomal degradative system. Ageing research reviews. 2009;8(3):199–213.

4. Aas SN, Hamarsland H, Cumming KT, et al. The impact of age and frailty on skeletal muscle autophagy markers and specific strength: A cross-sectional comparison. Exp Gerontol. 2019;125:110687.

5. Arribat Y, Broskey NT, Greggio C, et al. Distinct patterns of skeletal muscle mitochondria fusion, fission and mitophagy upon duration of exercise training. Acta physiologica (Oxford, England). 2019;225(2):e13179.

6. Balan E, Schwalm C, Naslain D, Nielens H, Francaux M, Deldicque L. Regular Endurance Exercise Promotes Fission, Mitophagy, and Oxidative Phosphorylation in Human Skeletal Muscle Independently of Age. Frontiers in Physiology. 2019;10:1088.

7. Carnio S, LoVerso F, Baraibar MA, et al. Autophagy impairment in muscle induces neuromuscular junction degeneration and precocious aging. Cell reports. 2014;8(5):1509–1521.

8. Wohlgemuth SE, Lees HA, Marzetti E, et al. An exploratory analysis of the effects of a weight loss plus exercise program on cellular quality control mechanisms in older overweight women. Rejuvenation research. 2011;14(3):315–324.

9. Bujak AL, Crane JD, Lally JS, et al. AMPK activation of muscle autophagy prevents fasting-induced hypoglycemia and myopathy during aging. Cell metabolism. 2015;21(6):883–890.

10. Carter HN, Kim Y, Erlich AT, Zarrin-Khat D, Hood DA. Autophagy and mitophagy flux in young and aged skeletal muscle following chronic contractile activity. J Physiol. 2018;596(16):3567–3584.

11. Wohlgemuth SE, Seo AY, Marzetti E, Lees HA, Leeuwenburgh C. Skeletal muscle autophagy and apoptosis during aging: effects of calorie restriction and life-long exercise. Exp Gerontol. 2010;45(2):138–148.

12. Zhou J, Chong SY, Lim A, et al. Changes in macroautophagy, chaperone-mediated autophagy, and mitochondrial metabolism in murine skeletal and cardiac muscle during aging. Aging. 2017;9(2):583–599.

13. Baumann CW, Kwak D, Liu HM, Thompson LV. Age-induced oxidative stress: how does it influence skeletal muscle quantity and quality? J Appl Physiol (1985). 2016;121(5):1047–1052.

14. Calvani R, Joseph AM, Adhihetty PJ, et al. Mitochondrial pathways in sarcopenia of aging and disuse muscle atrophy. Biological chemistry. 2013;394(3):393–414.

15. Green DR, Galluzzi L, Kroemer G. Mitochondria and the autophagy-inflammation-cell death axis in organismal aging. Science (New York, NY). 2011;333(6046):1109–1112.

16. Ziegler DV, Wiley CD, Velarde MC. Mitochondrial effectors of cellular senescence: beyond the free radical theory of aging. Aging cell. 2015;14(1):1–7.

17. Egan DF, Shackelford DB, Mihaylova MM, et al. Phosphorylation of ULK1 (hATG1) by AMP-activated protein kinase connects energy sensing to mitophagy. Science (New York, NY). 2011;331(6016):456–461.

18. Kundu M, Lindsten T, Yang CY, et al. Ulk1 plays a critical role in the autophagic clearance of mitochondria and ribosomes during reticulocyte maturation. Blood. 2008;112(4):1493–1502.

19. Laker RC, Drake JC, Wilson RJ, et al. Ampk phosphorylation of Ulk1 is required for targeting of mitochondria to lysosomes in exercise-induced mitophagy. Nature communications. 2017;8(1):548.

20. Wang C, Wang H, Zhang D, et al. Phosphorylation of ULK1 affects autophagosome fusion and links chaperone-mediated autophagy to macroautophagy. Nature communications. 2018;9(1):3492.

21. Nichenko AS, Southern WM, Tehrani KF, et al. Mitochondrial-specific autophagy linked to mitochondrial dysfunction following traumatic freeze injury in mice. Am J Physiol Cell Physiol. 2020;318(2):C242–c252.

22. Gheller BJ, Blum J, Soueid-Baumgarten S, Bender E, Cosgrove BD, Thalacker-Mercer A. Isolation, Culture, Characterization, and Differentiation of Human Muscle Progenitor Cells from the Skeletal Muscle Biopsy Procedure. Journal of visualized experiments : JoVE. 2019(150).

23. Klionsky DJ, Abdelmohsen K, Abe A, et al. Guidelines for the use and interpretation of assays for monitoring autophagy (3rd edition). Autophagy. 2016;12(1):1–222.

24. Turturro A, Witt WW, Lewis S, Hass BS, Lipman RD, Hart RW. Growth curves and survival characteristics of the animals used in the Biomarkers of Aging Program. The journals of gerontology Series A, Biological sciences and medical sciences. 1999;54(11):B492–501.

25. Nichenko AS, Southern WM, Atuan M, et al. Mitochondrial maintenance via autophagy contributes to functional skeletal muscle regeneration and remodeling. Am J Physiol Cell Physiol. 2016;311(2):C190–200.

26. Masser DR, Clark NW, Van Remmen H, Freeman WM. Loss of the antioxidant enzyme CuZnSOD (Sod1) mimics an age-related increase in absolute mitochondrial DNA copy number in the skeletal muscle. Age (Dordrecht, Netherlands). 2016;38(4):323–333.

27. Moulis M, Vindis C. Methods for Measuring Autophagy in Mice. Cells. 2017;6(2).

28. Smith NT, Soriano-Arroquia A, Goljanek-Whysall K, Jackson MJ, McDonagh B. Redox responses are preserved across muscle fibres with differential susceptibility to aging. Journal of proteomics. 2018;177:112–123.

29. Greising SM, Stowe JM, Sieck GC, Mantilla CB. Role of TrkB kinase activity in aging diaphragm neuromuscular junctions. Exp Gerontol. 2015;72:184–191.

30. Crupi AN, Nunnelee JS, Taylor DJ, et al. Oxidative muscles have better mitochondrial homeostasis than glycolytic muscles throughout life and maintain mitochondrial function during aging. Aging. 2018;10(11):3327–3352.

31. Kim J, Kundu M, Viollet B, Guan KL. AMPK and mTOR regulate autophagy through direct phosphorylation of Ulk1. Nature cell biology. 2011;13(2):132–141.

32. Call JA, Nichenko AS. Autophagy: an essential but limited cellular process for timely skeletal muscle recovery from injury. Autophagy. 2020:1–4.

33. Fuqua JD, Mere CP, Kronemberger A, et al. ULK2 is essential for degradation of ubiquitinated protein aggregates and homeostasis in skeletal muscle. FASEB journal : official publication of the Federation of American Societies for Experimental Biology. 2019;33(11):11735–11745.

34. Gallegly JC, Turesky NA, Strotman BA, Gurley CM, Peterson CA, Dupont-Versteegden EE. Satellite cell regulation of muscle mass is altered at old age. J Appl Physiol (1985). 2004;97(3):1082–1090.

35. Li Y, Lee Y, Thompson WJ. Changes in aging mouse neuromuscular junctions are explained by degeneration and regeneration of muscle fiber segments at the synapse. The Journal of neuroscience : the official journal of the Society for Neuroscience. 2011;31(42):14910–14919.

36. Pawlikowski B, Pulliam C, Betta ND, Kardon G, Olwin BB. Pervasive satellite cell contribution to uninjured adult muscle fibers. Skeletal muscle. 2015;5:42.

37. Valdez G, Tapia JC, Kang H, et al. Attenuation of age-related changes in mouse neuromuscular synapses by caloric restriction and exercise. Proceedings of the National Academy of Sciences of the United States of America. 2010;107(33):14863–14868.

38. Ribas V, Drew BG, Zhou Z, et al. Skeletal muscle action of estrogen receptor alpha is critical for the maintenance of mitochondrial function and metabolic homeostasis in females. Science translational medicine. 2016;8(334):334ra354.

